# Metabolic modeling of *Hermetia illucens* larvae resource allocation for high-value fatty acid production

**DOI:** 10.1101/2022.10.25.513765

**Authors:** Kristina Grausa, Shahida A Siddiqui, Norbert Lameyer, Karin Wiesotzki, Sergiy Smetana, Agris Pentjuss

**Affiliations:** Department of Computer Systems, Latvia University of Life Sciences and Technologies, Jelgava, Latvia; Technical University of Munich Campus Straubing for Biotechnology and Sustainability, Straubing, Germany; Institute of Microbiology and Biotechnology, University of Latvia, Riga, Latvia; German Institute of Food Technologies Quakenbrück, Germany

**Keywords:** Metabolic modeling, Insects, Fatty acids, *Hermetia illucens*, Resource allocation

## Abstract

All plant and animal kingdom organisms use highly connected biochemical networks to facilitate sustaining, proliferation and growth functions. While biochemical network details are well known, the understanding of intense regulation principles is still limited.

We chose to investigate *Hermetia illucens* fly at the larval stage as it is crucial for successful resource accumulation and allocation for the consequential organism’s developmental stages. We combined the iterative wet lab experiments and innovative metabolic modeling design approaches, to simulate and explain the *H. illucens* larval stage resource allocation processes and biotechnology potential. We performed time-based growth and high-value chemical compound accumulation wet lab chemical analysis experiments in larvae and Gainesville diet composition. To predict diet-based alterations on fatty acid allocation potential, we built and validated the first *H. illucens* medium-size stoichiometric metabolic model.

Using optimization methods like Flux balance and Flux variability analysis on the novel insect metabolic model, it predicted that doubled essential amino acid consumption increased the growth rate by 32%, but pure glucose consumption had no positive impact on growth. In the case of doubled pure valine consumption, the model predicted a 2% higher growth rate. In this study, we describe a new framework to research the impact of dietary alterations on the metabolism of multi-cellular organisms at different developmental stages for improved, sustainable and directed high-value chemicals.

**Significance Statement:** Metabolic modeling serves as a platform for researchers to investigate and study in depth the possible states of the system based on the existing knowledgebase (e.g. metabolic reactions, substrates, products and their stoichiometry). These models can be applied for different industrial applications, to simulate resource allocation potential and growth conditions. Moreover, these models predict the required diet for living organisms and insects to improve survival and growth rates and accumulate higher-value products, like fatty acids.

## Introduction

One of the major focal points in biology are processes which impact and shape organisms, giving rise to an enormous variation in different phenotypes including metabolism and even biomass composition in different growth and dietary conditions (1), (2), (3). In recent years, a consensus has been reached that resource allocation strategies and mechanisms are key points in determining an organism’s phenotype specificities (4), (5)

*Hermetia illucens* (Balck soldier fly, BSF) is widely recognized in the industry due to its ability to rapidly recycle organic waste and decompose the material into high-value lipids and proteins (6). BSF during the larval stage rapidly accumulates nutrients from the environment and increases its body weight according to sigmoidal growth function (7). BSF metabolizes organic waste into high-value chemicals without additional pre-processing, thus outperforming algae and other microorganism cultivation, purification and biotechnology processes and equipment costs. Thereby BSF has gained significant interest in food and feed, biofuel, pharmaceutical, lubricants and fertilizer sciences and industrial applications. The unbalanced fatty acid content of BSF larvae, however, is one of the main challenges for their industrial application (8), (9) and it warrants further investigation regarding the dietary impact on larvae body fat composition and growth rate to estimate their potential use for upcycling different agricultural waste products and residues.

Holometabolous insects go through several developmental stages, where each has a different objective and consequent functions in their life history. At each BSF life stage stochastic environmental conditions strongly impact nutrient and energy resource allocation metabolism (9). Insects possess different metabolic regulation mechanisms for (I) larvae, adults and (II) eggs, pupae, and diapausing individuals’ stages. While some aspects of resource allocation have been more extensively investigated (10), the mechanisms and expression of resource compensation on full genome-scale metabolism, especially at the first insect’s developmental stages, are still poorly understood (11). During larval development, BSF receives nutrients from the environment which are incorporated into large body mass or fat body, storing high-value acids.

Metabolic models successfully describe genotype-phenotype relationships for prokaryotic, eukaryotic, unicellular, and multi-tissue organisms, for example, Escherichia coli (12) and Saccharomyces cerevisiae (13). Metabolic modeling involves the analysis of an organism’s phenotypic responses (14) and predicts the carbon distribution and environmental perturbation impact on the metabolism (15), (16). Genome-scale metabolic modeling (GSM) has led to significant breakthroughs in health and systems medicine fields (17), in new nutrient uptake innovations (17), different novel software development (18), (19), new biotechnology applications (20), (21) ,(22) and other improvements in life science fields (23). Genome-scale metabolic models were also used to find new bio-based technologies to decrease waste streams and pollution (24), fine-tune genetic engineering design sets and explain an organism’s metabolic processes in different environments (24).

There already exist several insect models like central nervous system energy metabolism (25), the cost of metabolic interaction between insects and bacteria (26), evolutionary trajectories for bacterial and insect symbionts (27) and many more. However, up until now, there is no published *H. illucens in-silico* mathematical metabolic model, which could describe different resource allocation processes based on Gene-Protein-Reaction association, genetics, feed diet and environmental changes.

These models can fill existing knowledge gaps relating to phenotype-genotype relationships and biochemical network connectivity by combining in vivo experimental and publicly available omics datasets. Thus, we developed the first high-quality manually curated medium-scale BSF larvae metabolic model Hermetia_01 and applied it for modeling high-value fatty acid production and accumulation. The model incorporates experimentally measured dietary nutrient data (amino acids, monosaccharides) uptake rates and the accumulated biomass macromolecular composition. The model is adjusted for simulating accumulation, storage and resource allocation of the most abundant *H. illucens* fatty acids: lauric acid (C12:0), myristic acid (C14:0), palmitic acid (C16:0), palmitoleic acid (C16:1), stearic acid (C18:0), oleic acid (C18:1) and linoleic acid (C18:2).

Humans and other mammals are heterotrophs which need to consume other organisms to accumulate nutrients which cannot be synthesized within the body but are crucial for metabolic processes and health. Insect larvae present a great potential to combat the spreading shortages of essential nutrients, especially fatty acids for food and feed (28).

Saturated fatty acids (SFAs) have no double bonds in the carbon chain and they can be derived from animal fats as well as plant oils. Common dietary fatty acids include C12:0, C14:0, C16:0 and C18:0. In the past a diet high in SFAs had been associated with obesity, inflammation and negative cardiovascular outcomes (28), (29). However, more recent studies have shown no evidence of causation between a diet high in SFA and cardiovascular disease (30), (31), (32). In fact, some researchers report beneficial, cardioprotective, antioxidant, antimicrobial, antibiofilm and anti-viral and anti-cancer properties of some SFAs like C12:0 and C14:0 and they have even been shown to promote mitochondrial regulation in insulin-resistant macrophages (33), (34), (35), suggesting that the link between SFAs, cholesterol and heart disease is more complex than previously thought. C12:0 content in BSF larvae biomass was 29% of the total fatty acid profile, which makes it the most abundantly found fatty acid in *H. illucens* larvae according to our chemical analysis.

Monounsaturated fatty acids (MUFAs) are considered healthy dietary fats and are found in olive oil, avocado oil, nuts, seeds and some animal meat products. A diet rich in MUFAs is associated with a reduction in cancer, heart disease, inflammation and insulin resistance, (35), (35). C18:1 is capable of modulating the expression of oncogenes HER2, FASN and PEA3 (36), (37). The most abundant source of MUFAs in the human diet (~90% of total MUFAs) is C18:1 (38) which is an omega-9 fatty acid considered partially essential since endogenous synthesis can compensate for dietary deficiencies in organs such as the brain while leading to a reduction in concentration in other organs (39). C18:1 is also used in the pharmaceutical industry as an emulsifying or solubilizing agent (40). C18:1 content in BSF larvae biomass was 21% of the total fatty acid profile based on our chemical analysis.

Polyunsaturated fatty acids (PUFAs) are important parts of phospholipids, which have an essential impact on cell membrane structure and function (41). Higher animals like humans and other mammals cannot introduce double bonds in fatty acid chains beyond the 9th carbon atom, thereby rendering them incapable of synthesizing polyunsaturated fatty acids (PUFAs) (42). Since these fatty acids can only be absorbed through the diet, they are called essential fatty acids. and they include C18:2 belongs to PUFAs and is an essential omega-6 fatty acid mostly found in plant oils. C18:2 is a structural cell membrane component and a precursor for linolenic, arachidonic and other essential fatty acids involved in signaling molecule (hormone) production in animals (43) (44), thus should be supplied through diet in appropriate amounts. C18:2 content in BSF larvae biomass was 20% of the total fatty acid profile based on our chemical analysis.

We used the novel metabolic model Hermetia_01 for the execution of two sets of *in-silico* BSF larva metabolism experiments **(1)** predicting specific growth rate changes based on dietary alterations (under conditions of doubled glucose or doubled essential amino acid uptake); and **(2)** predicting fatty acid (C12:0, C14:0, C16:0, C16:1, C18:0, C:18:1, C18:2) synthesis rate changes with doubled glucose uptake and calculating dietary carbon conversion efficiency. The first set of experiments indicated that only an increase in essential amino acid uptake rates resulted in a higher specific growth rate of BSF larvae, while additional glucose did not affect the growth rate. The second set of experiments increased all above listed fatty acid synthesis rates when glucose consumption was doubled. However, the dietary carbon conversion efficiency was lower for fatty acid synthesis in the conditions of doubled glucose uptake, in contrast to our reference dietary experimental data.

## Results

### The first medium-scale metabolic model for *Hermetia illucens*

Considering that no metabolic model of *Hermetia illucens* larva has previously been published despite the growing interest in its ability to rapidly recycle organic waste and decompose the material into high-value lipids and proteins (44), (45), (9), we generated and manually curated Hermetia_01, which describes the metabolism of *H*. *illucens* at the larval development stage. Our model covers a total of 326 metabolites, 407 reactions, 471 genes and 3 compartments and provides means to study in-depth and examine biochemical pathways involved in fatty acid and amino acid synthesis by BSF larvae to assess their potential as a sustainable source of lipids and proteins not only for food and feed but also for pharmaceutical, industrial and even thermal energy storage and industrial exhaust heat utilization and recycling applications (46) (47), (48). The unbalanced fatty acid content of BSF larvae is one of the main challenges for their use in food and feed (9) and warrants further investigation regarding the dietary impact on larvae body fat composition. For this purpose, we focused on lipid metabolism and specific fatty acid (C12:0, C14:0, C16:0, C18:0, C18:1, C18:2) synthesis and accumulation in the biomass during the development of the metabolic model. The 6 previously mentioned fatty acids were selected since taken together they represent ~90% of the total fatty acid composition of BSF larvae based on our chemical analysis. Hermetia_01 has been tailored for modeling, predicting and optimizing fatty acid and amino acid metabolism processes and resource allocation using different diets and substrates.

### Biomass Composition and Reaction

The given diet, substrate composition and environmental factors affect the growth and development of any organism resulting in varying biomass composition specific to that environment. Some fatty acids, mostly saturated (C12:0, C14:0, C16:1) were shown to be synthesized by BSF larvae and not bioaccumulated (47), (49).

Using our chemical analysis experiments of amino acid, fatty acid and macromolecular composition of *H. illucens* larvae (Fig. 1A), we formulated a basic level biomass objective function (50) which describes the composition of the cell and energetic requirements necessary to generate biomass content from metabolic precursors and sustain cell proliferation. Our biomass objective function (Fig. 1B) consists of amino acids, fat (triglyceride), starch (glycogen), sugars (mono- and disaccharides) and glycerol with respective ratios in mmol per one dry weight (DW) larva calculated based on our experimental measurements and cholesterol which was not captured during chemical analysis but is a major constituent of animal cell membranes and therefore was approximated relative to other biomass macromolecules using previously published *Drosophila* metabolic network (Schönborn et al., 2019). We normalized our DW biomass macromolecule ratios to 100%.

**Figure 1.**
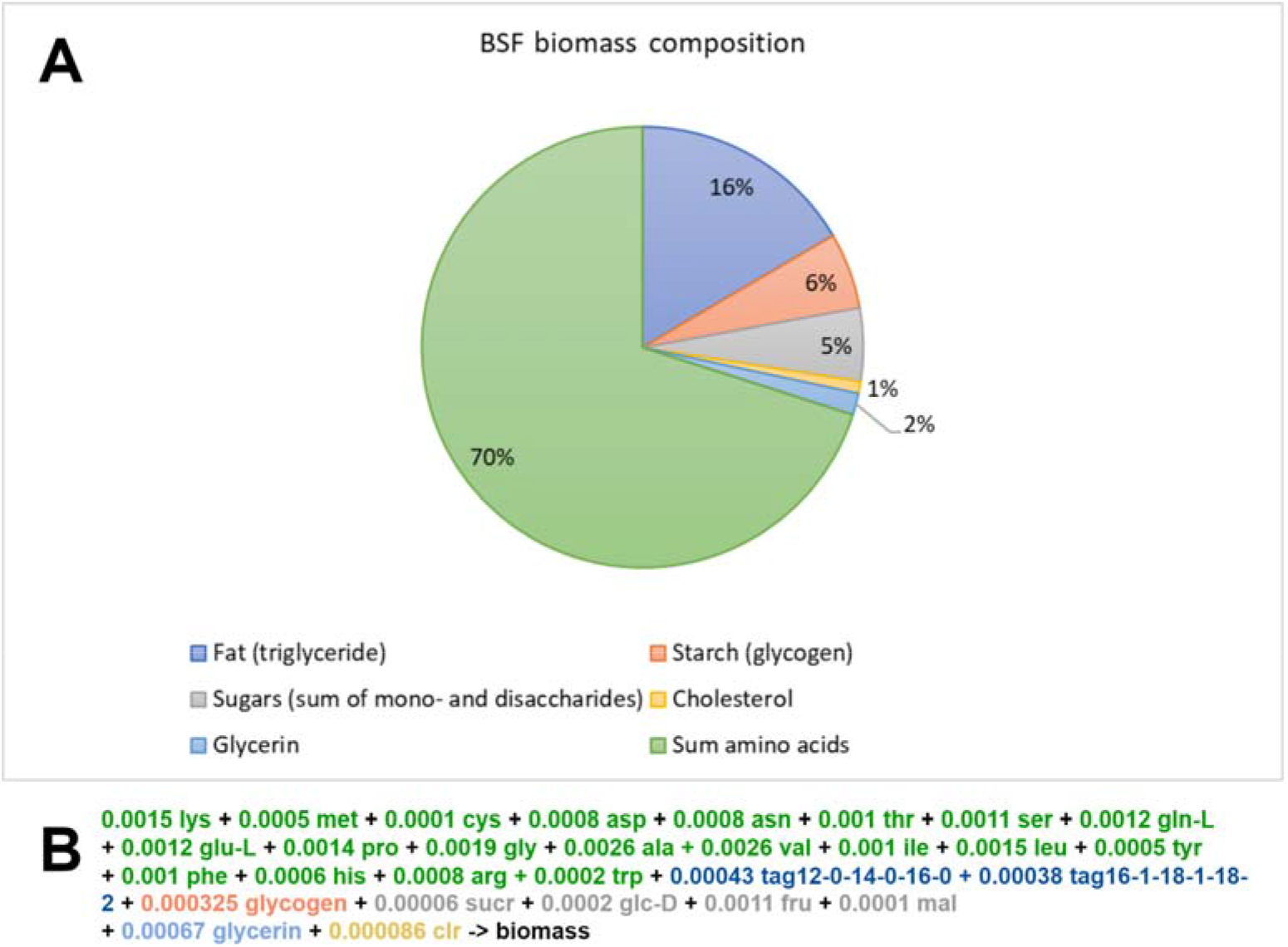
Biomass composition and function of BSF larva. (A) Normalized DW biomass metabolite ratios of BSF larva. (B) Biomass objective function based on the chemical analysis experiments.

Triglycerides (TAG) are esters derived from glycerol and three fatty acid chains of varying lengths and compositions. These esters are the main component of animal fat which are stored in cellular organelles called lipid droplets or adiposomes (51) inside specialized storage organs like adipose tissue in mammals or the fat body in insects (52). In the case of holometabolous insects, such as BSF, fat storage is essential during the larval development stage since triglycerides serve as energy storage and resource (53). For modeling purposes, we first estimated the most abundant fatty acids in BSF larvae based on our chemical measurements of fatty acid composition (Table 1), which were lauric acid (C12:0) 29%, myristic acid (C14:0) 6%, palmitic acid (C16:0) 10%, palmitoleic acid (C16:1) 6%, oleic acid (C18:1) 21% and linoleic acid (C18:2) 20%. Taken together these fatty acids make up 90% of the total fatty acid composition in BSF larvae (Supplementary file 10) (Table 1). Our chemical measurements of larvae fatty acid composition indicate 49% of saturated fatty acids (C12:0, C14:0, C16:0, C18:0) and 48% of unsaturated fatty acids (C14:1, C16:1, C18:1, C18:2) of which 28% were monounsaturated (C14:1, C16:1, C18:1) and 20% polyunsaturated fatty acids (C18:2). Consequently, we extended our metabolic model by incorporating diacylglycerol and triacylglycerol biosynthesis pathway (TRIGLSYN-PWY) for TAG (12:0/14:0/16:0) and TAG (16:1/18:1/18:2) synthesis which we used in biomass objective function formulation for BSF larva with ratio 49:51 of total biomass fat respectively (Fig. 1A).

**Table 1.**
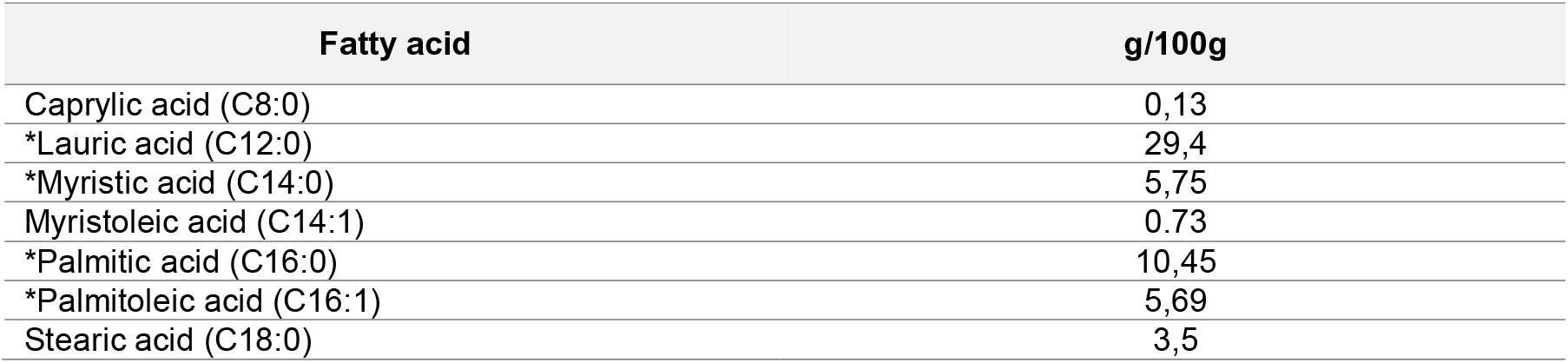
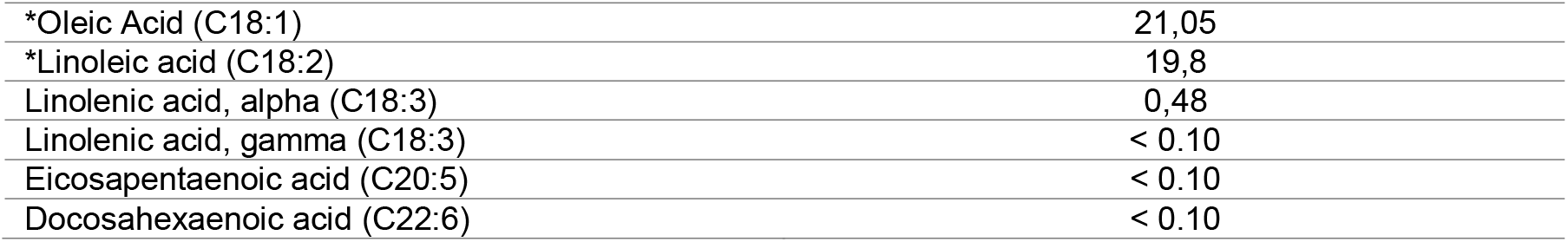
Fatty acid chemical analysis results of *H. illucens* larva. (* notify amino acids which included in Hermetia_01 metabolic model biomass composition function).

### Model validation and evaluation

The steady-state assumption is frequently used in metabolic modeling and serves as a means to validate the interconnectivity of a biochemical network and the feasibility of its parameters. This assumption predicts that internal metabolite production and consumption must be balanced (54) meaning there is no accumulation or deficit of metabolites. Many metabolic modeling analysis methods incorporate the steady-state assumption in their formulation, one such method is flux balance analysis (FBA) (55) which is a constraint-based linear programming method dependent on constraints enforced by the metabolic network stoichiometry and is the most frequently used method for metabolic network analysis. We began the evaluation of our metabolic network by validating biomass precursor production, defining transport reactions for each biomass component and performing FBA on each of these reactions as the optimization objective. This allowed us to ensure that each biomass component is capable of carrying flux (being produced) and is not blocked. We also used the FBA method as a means to validate the feasibility of our metabolic network and the applied constraints based on our chemical measurements. We continued the validation process by performing FBA on biomass function as the optimization objective and having transport reactions constrained according to the given diet during wet lab experiments. This allowed us to extract the maximal theoretical biomass growth rate. Unfortunately, the actual growth rate was not captured during wet lab experiments.

Next, we evaluated our metabolic model based on several parameters described in a previously published protocol for generating a high-quality genome-scale metabolic reconstruction (51). We performed Cobra Toolbox functions on our metabolic network in the MATLAB environment and estimated 2.45% dead-end metabolites (*detectDeadEnds*), 16.71% blocked reactions (*findBlockedReaction*), 19.66% unbalanced reactions (*checkBalance*) and 18.43% exchange reactions (*findExcRxns*) within our model and compared the results with other metabolic networks (see section Metabolic network comparison).

We also validated our DW biomass fat (16%) and amino acid (70%) ratios of BSF larva which were in line with previous literature (6). Fatty acid profile measurements consisting of lauric acid (C12:0) 29%, myristic acid (C14:0) 6%, palmitic acid (C16:0) 10%, palmitoleic acid (C16:1) 6%, oleic acid (C18:1) 21% and linoleic acid (C18:2) 20% were within the exact range described in previous literature (7).

### Metabolic network comparison

The result metabolic model of *H. illucens* at the larval development stage covers not only the central metabolism pathways (glycolysis, pentose phosphate pathway, TCA) but also amino acid and lipid metabolism pathways since BSF larva are widely recognized for their high lipid and protein content, (9). The metabolic model covers 326 metabolites, 407 reactions, 471 genes and 3 compartments (cytosol, mitochondria, extracellular) in total. We compared our metabolic network of BSF larvae with other curated and publicly available metabolic networks including E. coli central and genome-scale metabolism, S. cerevisiae and Human (Table 2). Relative to other metabolic networks Hermetia_01 excels with a low percentage (2.45%) of dead-end metabolites, suggesting sufficient interconnectivity of the biochemical network where only a few metabolites participate in just one reaction or can only be consumed or produced. Blocked reactions which are incapable of carrying a flux account for 16.71% of our metabolic network. Reactions unbalanced elementally (19.66% in Hermetia_01) include exchange reactions (18.43% in Hermetia_01) and reactions synthesizing or consuming metabolites with no charged chemical formula provided, in our case such metabolites were biomass macromolecules used to split and simplify the biomass reaction formulation and protein class metabolites (ferricytochrome b5, ferrocytochrome b5) used for C18:1 and C18:2 synthesis.

**Table 2.**
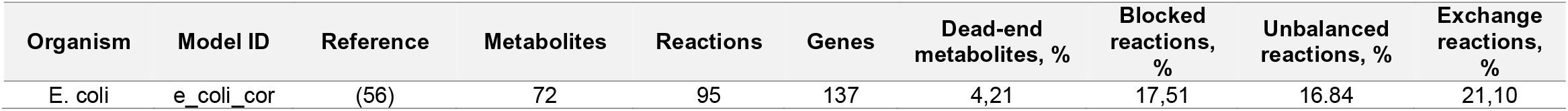
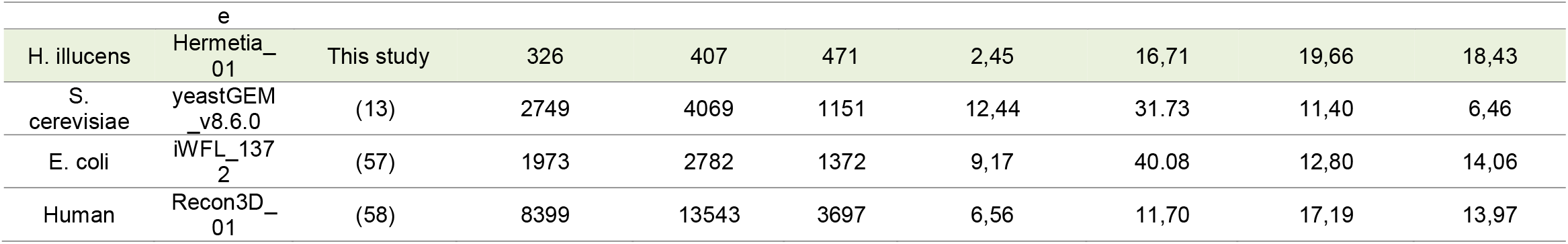
Comparison of Hermetia_01 to other curated and publicly available metabolic networks.

### Metabolic Flux Potential as Predicted by Flux Variability Analysis/ Resource allocation

To assess differences in resource allocation mechanisms within the Hermetia_01 metabolic network of BSF larvae based on the given diet, we performed several *in-silico* experiments utilizing flux variability (FVA) (59) analysis provided by the Cobra Toolbox software. The FVA method calculates maximal and minimal biochemical network reaction rate values for each reaction in a metabolic network while maintaining (in our case) 90% of the maximal biomass-specific growth rate per day as described by (60).

We performed two sets of *in-silico* experiments **(1)** simulation of biomass growth rate changes based on our dietary data and under conditions of doubled glucose (2x_glc) or doubled essential amino acid (2xEAA) uptake rates; **(2)** simulation of different fatty acid (C12:0, C14:0, C16:0, C16:1, C18:0, C:18:1, C18:2) synthesis rate changes with doubled glucose consumption and a calculation of dietary carbon conversion efficiency.

In growth rate change experiments, we defined 3 different design sets to obtain the theoretical substrate carbon distribution potential and predict substrate (glucose and amino acids) uptake change impact on the organism's specific growth rate. **(1)** Doubling glucose consumption rate in the Hermetia_01 medium-size metabolic model did not show any increase in specific growth rate when compared to a dietary unchanged state based on our experimental data (Fig. 2) for the BSF larvae, similar to the predictions of the *FlySilico* model (61) of *Drosophila* larvae.

**Figure 2.**
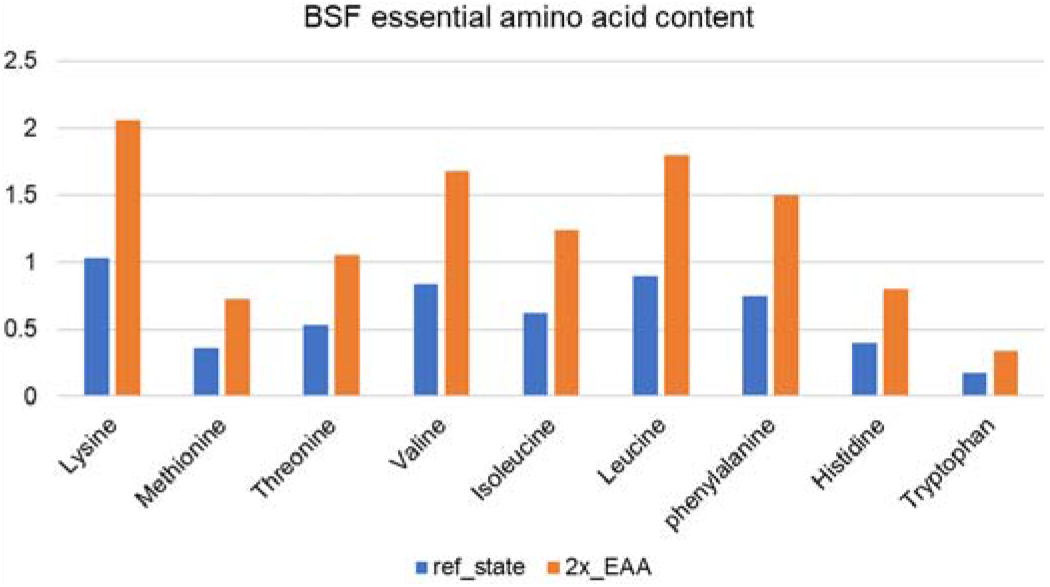
*H. illucens* larva EAA composition, g/100 g larvae.

Mammalians are heterotrophic organisms which require the consumption of specific nutrition through diet, like EAA and others. We prepared an optimization design set, where **(2)** we doubled all EAA consumption rates (Fig. 2)

The Hermetia_01 showed a theoretical potential to increase BSF specific growth rate by 32% (Fig. 3). The model could not simulate all amino acid degradation metabolism processes, but the **(3)** doubled Valine consumption and later degradation in chemical building blocks, resulting in a 2% increase in specific growth rate.

**Figure 3.**
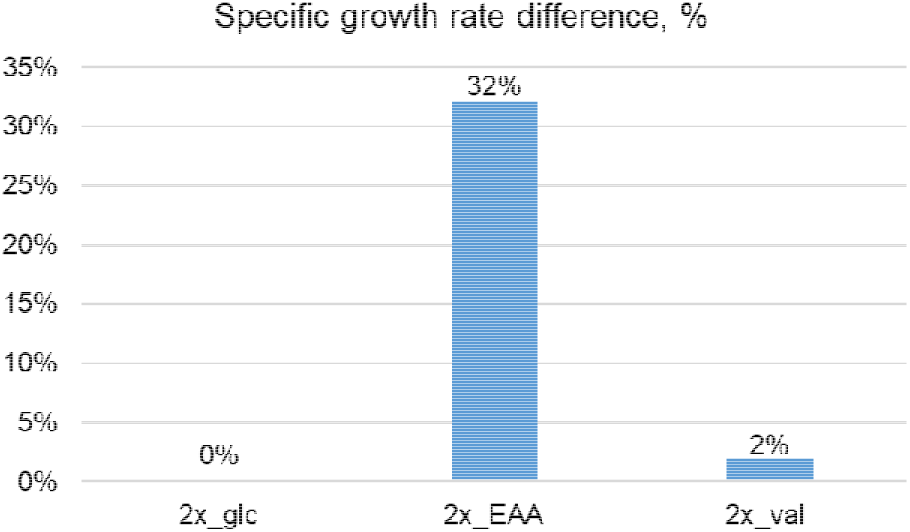
Predicted growth rate of BSF larvae for the doubled glucose uptake state (2x glc), the doubled essential amino acid uptake state (2x EAA) and the doubled valine uptake state (2x val). Comparisons are made based on our dietary experimental data in the Hermetia_01model and changes calculated in percent.

Fatty acid (C12:0, C14:0, C16:0, C16:1, C18:0, C18:1, C18:2) synthesis increased by doubling glucose consumption rate in the Hermetia_01 (Fig. 4), especially linoleic acid (C18:2) which increased from 0.029 to 0.051 mmol/larva/day.

**Figure 4.**
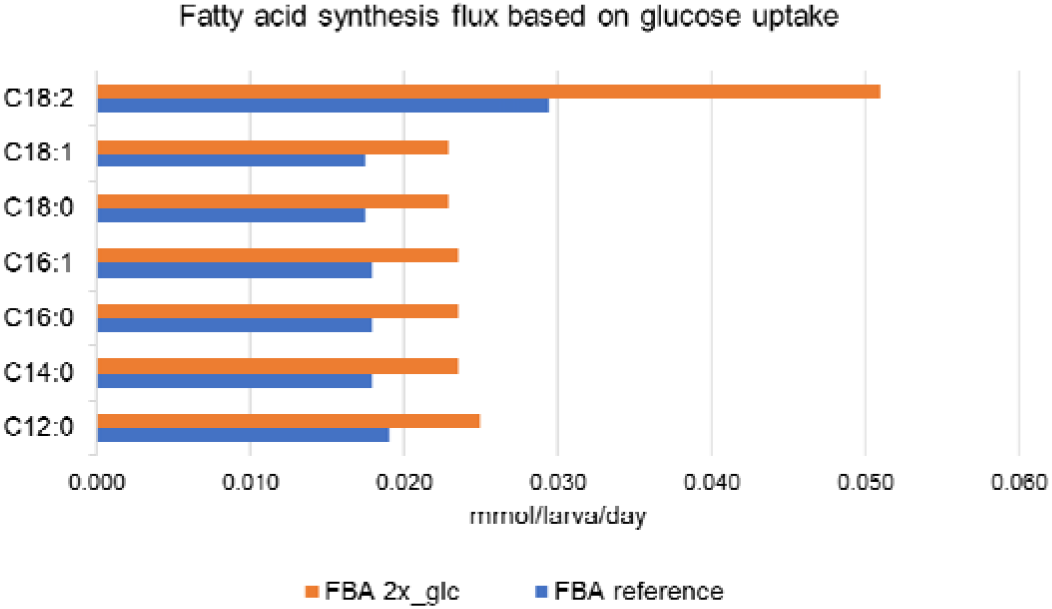
Fatty acid synthesis flux rates for the reference state (ref state) and doubled glucose uptake (2x glc). However, the conversion efficiency of dietary carbon to high-value fatty acids dropped for the doubled glucose state in comparison to the reference state. This indicated that the additional nutrition was not used sufficiently (Fig. 5) for fatty acid synthesis. One exception was linoleic acid (C18:2) which resulted in a nearly doubled production rate compared to the reference state (Fig. 5).

**Figure 5.**
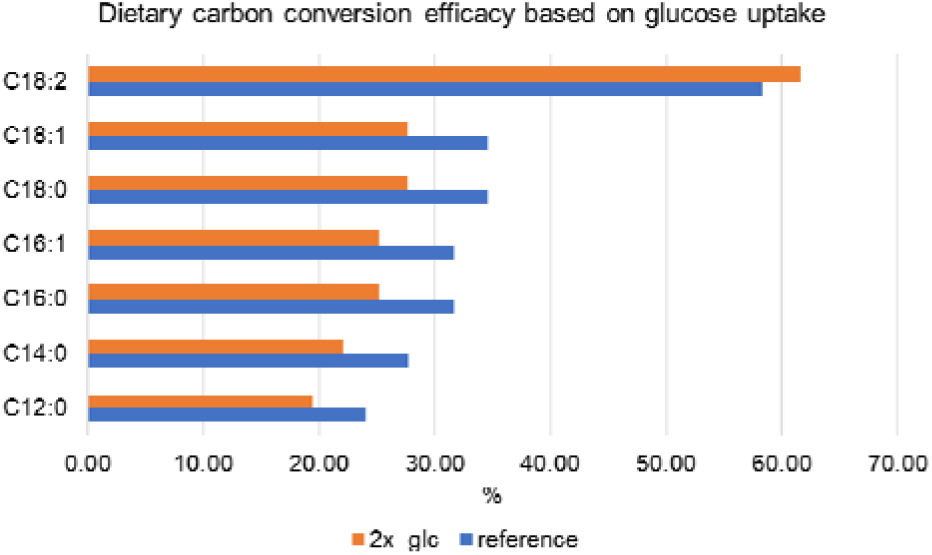
The dietary carbon conversion efficiency for the reference state (ref state) and under the conditions of doubled glucose uptake (2x glc).

The performed growth rate change *in-silico* experiments indicate that the low growth rate of BSF larvae is caused by amino acid deficiencies rather than the lack of energy resources, which is in line with the prognosis of other researchers studying the dietary impact *H. illucens* larva metabolism and growth (62). Therefore, to optimize BSF larva biomass production and additional protein supplementation in the diet should be considered. As for optimizing fatty acid (C12:0, C14:0, C16:0, C16:1, C18:0, C:18:1, C18:2) synthesis and accumulation in the BSF larvae biomass, an optimal glucose uptake threshold should be determined and tested during future wet lab experiments, considering the dietary carbon conversion efficiency changes when glucose uptake is increased. This knowledge can be applied to adjust the BSF larvae diet composition to increase the biotechnological potential of different product metabolites and for determining the best waste products or residues for BSF larvae to recycle.

## Discussion

### Potential fatty acids production in insects

Insect use in food and feed even though currently strictly regulated has attracted the attention of many researchers worldwide due to the rich protein and fat composition of many insects (61), (63). The potential insect utilization for agricultural waste recycling, reuse and supplementation in feed is particularly relevant for farmers, food and feed and potentially pharmaceutical manufacturers. There are several important advantages of insect rearing compared to livestock farming, which includes reduced resource requirements, high feed conversion efficiency and the potential for recycling and reuse of industrial side streams or byproducts, (63). Apart from the high-quality protein content necessary for feed, fatty acids serve as energy sources for heterotrophic organisms in the world. In this study, we focused on those fatty acids, which were most abundantly found in BFS composition (C12:0, C14:0, C16:0, C16:1, C18:0, C18:1, C18:2) and also have high economic value in food and feed (46), industrial, pharmaceutical industries and recycling, (47), (64). 49% of the total BSF larvae fat composition were saturated fatty acids (C12:0, C14:0, C16:0, C18:0) and 48% of unsaturated fatty acids (C14:1, C16:1, C18:1, C18:2) of which 28% were MUFAs (C14:1, C16:1, C18:1) and 20% PUFAs (C18:2).

We found that there was no previously published metabolic model of BSF, which could utilize genome sequence and gene-protein-reaction association (GPR). Hermetia_01 is the first metabolic model, which is focused on specific fatty acid production by implementing diet uptake and biomass composition. The biomass objective function is used in the model to describe necessary the organism’s proliferation resources.

We performed different BSF resource allocation potential simulations to analyze (a) specific growth and (b) fatty acid synthesis rates in larva fat body by doubling glucose or EAA uptake rates in contrast to our reference experimental dietary data.

Since BSF is a multi-tissue and multi-compartmented organism, the proposed metabolism simplification by model Hermetia_01 may not provide ideal optimization results, but still, we believe that fatty acid production from carbohydrates in most organisms is synthesized by central metabolic biochemical pathways via acetyl-COA. We introduced detailed fatty acid production pathways by integrating lumped reactions (65). These reactions represent all metabolites involved in the fatty acid synthesis metabolic pathway, have manually corrected stoichiometry and are mass balanced.

Comparing the *FlySilico* (65) *Drosophila melanogaster* with Hermetia_01 as insect-based metabolic models, Hermetia_01 contains a more detailed description of TAG synthesis by incorporating different saturated fatty acids (C12:0, C14:0, C16:0, C18:0) and unsaturated fatty acids (C14:1, C16:1, C18:1, C18:2) instead of a generalized TAG description, which lacks the specificity for lipid metabolism and resource allocation process modeling. Thus Hermetia_01 is capable of predicting the impact of dietary changes on biomass production as well as the fatty acid resource allocation processes in the fat body during the larval development stage.

Using gene-protein-reaction association properties and steady-state assumption the model can be used to predict the necessary diet composition to achieve a desirable insect biomass composition and specific growth rate (Fig.3) given access to quality experimental omics data. Model versions are available in SBML, MAT, XLSX and JSON formats, which allows widespread use of various paid and open-access metabolic modeling tools and environments. We have also provided the Hermetia_01 metabolic model biochemical pathway visual layout, which is compatible with an in-house built web application for interactive metabolic flux analysis and visualization tool IMFler (66). Experts can easily perform the FBA or FVA metabolic analysis algorithms in IMFLer utilizing the user-friendly interface with no need for installing metabolic modeling tools on a local computer.

The unbalanced fatty acid content of BSF larvae, however, is one of the main challenges for their industrial application (6). Hoc et. al. also indicates that fatty acids can be directly accumulated from diet and de novo synthesized in BSF (9), suggesting the need for further experiments on amino acids and the role of carbohydrate level in BSF diets. Our Hermetia_01 metabolic model already allows us not only to predict dietary carbon accumulation or resource allocation by de novo fatty acid biosynthesis but also shows the dietary EAA and carbohydrate impact on metabolism.

Hermetia_01 model can predict how dietary glucose uptake changes affect specific fatty acid synthesis in the fat body during the larval development stage. We observed a positive impact of an increased glucose uptake rate on fatty acid synthesis (Fig. 3), where optimization results showed a ~ 20 % increase in fatty acid by doubled glucose consumption. One exception was linoleic acid (C18:2) which resulted in a nearly 60 % increase.

Many experiments have been done, where the direct accumulation of fatty acids from the environment was observed, but not synthesized by de-novo biochemical pathways. The diet composition and rearing environmental conditions in most cases are not similar, thus some reports have contradicting results, where oleic and linoleic acid proportions can significantly vary (67) (68). Further investigation is necessary to better understand fatty acid accumulation processes.

Our Hermetia_01 metabolic model optimization results also suggest that BSF significant growth rate changes can be achieved by increasing the dietary protein content, especially EAA amounts (Fig. 2.), indicating that glucose or other mono sugar increase alone is not enough to improve the specific growth rate. Similar conclusions have been reported by other researchers studying the dietary impact on *H. illucens* larva metabolism and growth (62).

The novel Hermetia_01 metabolic model as well as our *in-silico* experiments can be applied for adjusting the BSF larvae diet composition to increase the biotechnological potential of different product metabolites and for determining the best waste products or residues for BSF larvae to recycle.

This is becoming especially fascinating nowadays as new human diets arise, like low carbohydrate and keto diets. Insect farming requires fewer resources, has the potential to extract, accumulate and reuse high-value nutrients from waste, and can be successfully employed in small-scale as well as large-scale production. Insect breeding requires less water on 1 kg compared to red meat and other livestock conventional practices. In this context insect metabolic modeling rapidly gains interest and demand in scientific, industrial food and feed, medical and other biotechnological industries and applications.

## Materials and Methods

The insect-rearing sector is rapidly growing in Europe and becoming one of the sustainable choices to produce feed, food and other substances. To ensure the efficiency and sustainability of insect production, they should rely on wastes and under-utilized side streams from agri-food systems (69), (70). At the same time, currently, there is no specific chemically defined diet, which can be used as a reference and basis for the cultivation of mass-produced insects. Current studies rely on the Gainesville Diet, which has an approximate chemical composition and can have variations for the specific chemicals in the scope of 20-30%. Multiple studies, using this diet as a reference get different, sometimes opposite results (71), (72).

Recent studies, determining the impact of specific nutrients on the development of *H. illucens*, indicated that they indeed have an impact and diets of insects would require high amounts of proteins (73) and lipids (74). However, they do not provide a detailed model for the optimal chemically defined diet. Such a diet would allow for improvements in experimental and theoretical studies, optimization of insect production and identification of suitable waste and side-stream mixes.

### Larvae rearing experiments

Newly hatched old 9 days larvae from eggs of the black soldier fly, *H. illucens* (Diptera: Stratiomyidae) were used for the experiments and reared on chicken feed (GS Gänse-Mastfutter 1, metabolizable energy content 12,1 MJ ME, 70 % moisture) for around 7 days.

They were then transferred to the substrate containing 10% glucose, 20% egg white protein and 70% water. The experiment was conducted in a plastic box (Transoplast, 60 cm x 40 cm x 12,5 cm), kept at 27° C and 60% humidity. In each box, 450g larvae were placed. The experiment was stopped after nine days of feeding. The substrate was weighed when applied, as was the total remaining material at the end of the experiment.

### Larvae wet weight measurements

For single larvae weight, the combined 30 larvae were recorded and divided by 30 to calculate the single larvae weight. The larvae were separated from their feed. At the end of the experiment larvae were collected, then placed in a sieve and thoroughly washed with water with extreme care, dried on a piece of paper, and finally weighed.

### Experimental chemical analytics and analysis methods

#### GC-MS measurements

All the studies were conducted with the help of Hewlett-Packard 5890 (GC)/5970 mass selective detector (MSD) in electron impact (EI) mode (70eV) with a configuration setup with a fused silica capillary column (HP-5ms, 30 m × 25 mm ID, 0.25 μm film thickness; Agilent J& W Scientific, Folsom, CA, USA). The 2 μL sample was inserted using splitless mode. The cover was kept at a temperature of 70 °C for 3 minutes. After that, it was increased to 280 °C at a rate of 25 °C/min and then kept at this temperature for 5 minutes. Helium was used as the transporting gas and was fed in at a constant rate of 1 mL/min. The injector temperature was kept constant at 240 °C and the terminal temperature was kept at 290 °C. The mass spectrum of the MCF-derivatized amino acids and inherent criteria were gathered in either full-scan (50-350 m/z) or selected ion monitoring (SIM) acquisition modes.

#### Fatty Acid Profile

In a modification, after freeze-drying, the samples are heated directly with a methanolic sulfuric acid solution. The resulting fatty acid methyl esters are then extracted using n-hexane and analyzed according to the standard procedure by capillary gas chromatography. Determination based on the ISO 12966-2 standard (section 5.5).

#### Sugars (Glucose, Fructose, Sucrose and Maltose) and Glycerol

After homogenizing, the samples were prepared by L05.00-10, Section 7.2.1 (Federal Office for Consumer Protection and Food Safety, Germany, official collection of examination procedures) (5g sample, Carrez clarification). The solutions obtained were measured using HPLC and a refractometric detector by the Method of the German Institute of Food Technologies

#### Starch

The method comprises two determinations. In the first, the sample is treated with dilute hydrochloric acid. After clarification and filtration, the optical rotation of the solution is measured by polarimetry. In the second, the sample is extracted with 40 % ethanol. After acidifying the filtrate with hydrochloric acid, clarifying and filtering, the optical rotation is measured as in the first determination. The difference between the two measurements, multiplied by a known factor, gives the starch content of the sample. Determination according to Appendix III L of Regulation (EC) No. 152/2009 laying down the methods of sampling and analysis for the official inspection of feed.

#### Fat

With the addition of an internal standard, fat from the samples was extracted and at the same time, alkaline digested. After extraction, the potassium salts are converted into free fatty acids and the proportion of fatty acids is determined by gas chromatography. The fat content, calculated as triglyceride content in a 100g sample, is calculated using response and conversion factors and a calibration standard fat. Determination according to AOAC Method PVM4: 1997 (Caviezel ®).

### Metabolic network reconstruction of *Hermetia illucens* larva

Insects are multi-tissue, multiorgan animals and metabolic modeling of such organisms can be challenging. Extensively detailed genome-scale metabolic networks, which describe whole-body systems (e.g. Human, *A. thaliana*), can offer new and profound insights into organism metabolism. However, the development of such models requires decades-long research, extensive literature and a knowledge base of the target organism, its organ systems, genetics and other details and specificities. A more simplified approach for modeling complex organisms, where the whole organism is described as one comprehensive organ, has previously been used by other researchers to model other similar insect species (61) and is sufficient enough to model metabolism and fatty acid production.

We generated a draft reconstruction based on genome-scale model development protocol using *H. illucens* genome annotation data (available on (12.10.2022): https://www.ncbi.nlm.nih.gov/assembly/GCF_905115235.1). A draft model was generated with RAVEN (75) metabolic modeling. Next, we applied manual reaction curation using biochemical databases like MetaCyc (76), BioCyc (76), KEGG (77), PubChem (35), ChEBI (78) and additionally used a previously published and curated Drosophila larval central metabolic model data for medium size central metabolism reference, focusing on pathways covering our quantitative experimental measurements and fatty acids optimization potential. Our metabolic network covers a total of 326 metabolites, 407 reactions, 471 genes, and 3 compartments and the model scale includes extended central metabolism pathways analogue to the functionality of E. coli central metabolism (79). The interactive metabolic model visualization was drawn by IMFLer online modeling tool (66) (Fig. 6).

**Figure 6.**
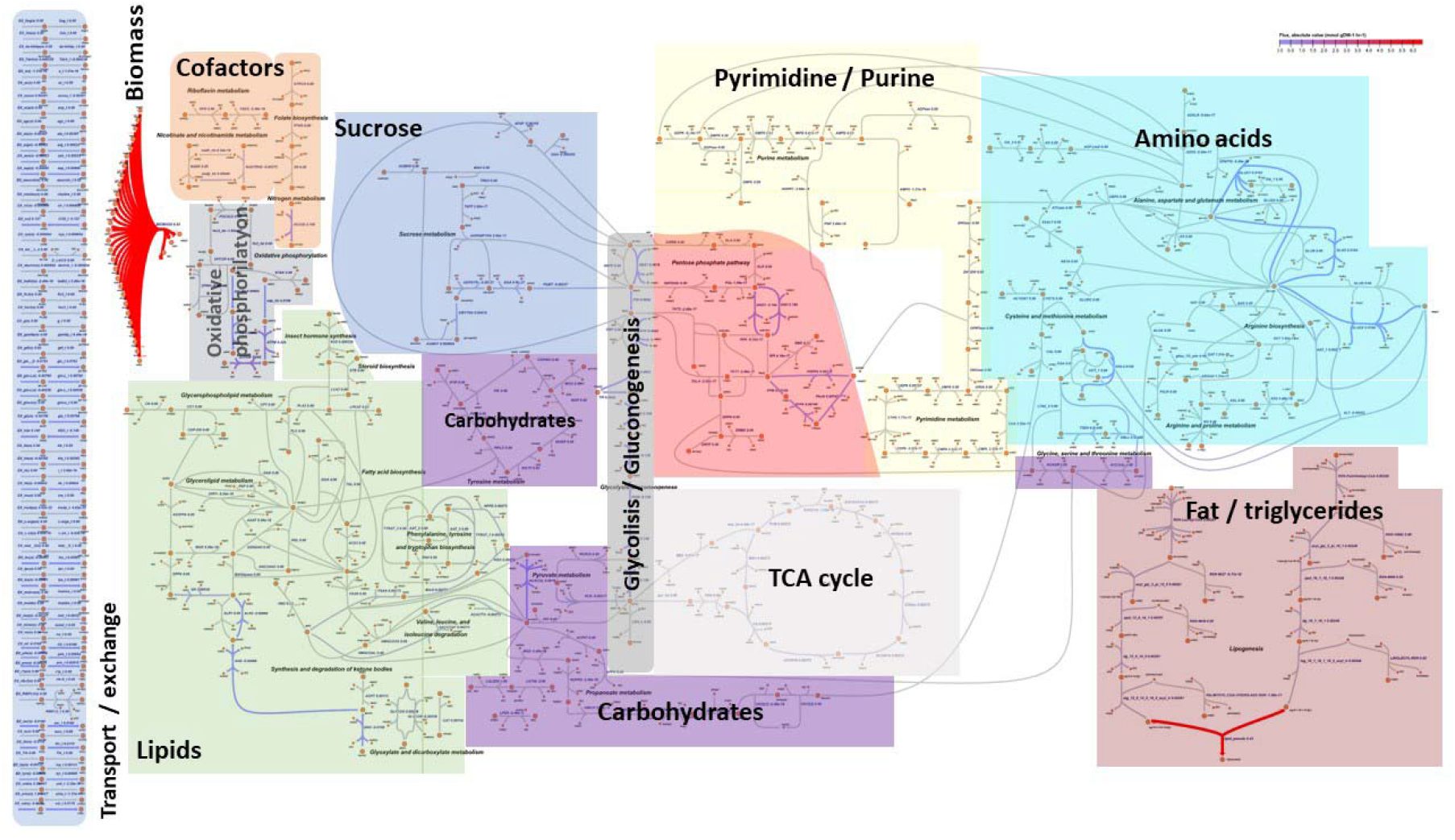
Metabolic model biochemical pathways of *H. illucens* for resource allocations analysis for high-value fatty acids accumulation. Each color represents a separate pathway. The model includes all necessary biochemical pathways to accumulate fatty acids (improved from (61)).

To recharge ferricytochrome b5 which is used for oleic and linoleic acid synthesis we included an additional reaction called cytochrome-b5 reductase reaction (BioCyc ID: CYTOCHROME-B5-REDUCTASE-RXN). Furthermore, we extended our metabolic model by incorporating diacylglycerol and triacylglycerol biosynthesis pathway (TRIGLSYN-PWY) according to the redGEM approach (65), by lamping together several reactions in a single one, to model fat storage of TAG (12:0/14:0/16:0) and TAG (16:1 or 18:0/18:1/18:2) within BSF larvae biomass.

The metabolic network was reconstructed using the MATLAB environment and optimization, analysis and validation tasks were done using Cobra Toolbox 3.0 software (80). We also integrated our experimental measurements on transport reactions representing the substrate in which larvae were grown and the given diet during wet lab experiments.

### The biomass metabolic reaction

We used quantitative BSF larvae chemical analysis experiments of amino acids, fat (triglyceride), starch (glycogen), sugars (mono- and disaccharides) and glycerol to calculate their respective ratios per one dry weight (DW) larva. Since cholesterol concentration was not captured during the wet lab experiments but is a major constituent of animal cell membranes, we approximated the ratio of cholesterol relative to other biomass metabolites using a previously defined and published Drosophila larva biomass function. Chemical analysis experiments of water, carbohydrates and ash were omitted from biomass composition and the rest of the biomass macromolecule ratios were normalized to 100% of the biomass. Biomass metabolite ratios (Fig. 1A) were then converted to mmol per one BSF larva DW and used in biomass reaction definition. We selected TAG (12:0/14:0/16:0) and TAG (16:1/18:1/18:2) in our biomass reaction formulation with a ratio of 49:51 of total biomass fat respectively. These triglycerides were chosen for the reason that they consist of fatty acids most abundant in BSF larvae according to our chemical analysis data of fatty acid composition and together account for 90% of total fatty acids.

### Lipid metabolism

We focused on lipid metabolism and specific fatty acid (C12:0, C14:0, C16:0, C16:1, C18:0, C18:1, C18:2, C18:3, C20:5, C22:6) synthesis while developing a metabolic model of BSF larva. These fatty acids were of special interest for modeling *H. illucens* larvae metabolism due to their high content in the organism captured by fatty acids profiling measurements or because of their biotechnological significance (omega-3 fatty acids: C18:3, C20:5, C22:6) and prospects for optimizing, analyzing and predicting their yields using bioconversion by recycling and therefore reducing industrial and agricultural waste (81).

Lipid metabolism reaction complexes included in our metabolic model cover several pathways:

1. Fatty acid biosynthesis initiation (type I), PWY-5966;
2. Palmitate biosynthesis I (type I fatty acid synthase), PWY-5994;
3. Tetradecanoate biosynthesis (mitochondria), PWY66-430;
4. Stearate biosynthesis I (animals), PWY-5972;
5. Oleate biosynthesis II (animals and fungi), PWY-5996;
6. Linoleate biosynthesis II (animals), PWY-6001;

The consecutive biochemical reactions listed in the pathways above were summed up into lumped reactions (substrates and product metabolites) in one reaction for each respective Acyl-CoA synthesis coupled with a hydrolase reaction to synthesize the respective fatty acid. Oleoyl-CoA and linoleoyl-CoA synthesis required a cytochrome-b5 reductase reaction (BioCyc ID: CYTOCHROME-B5-REDUCTASE-RXN) which was also added to the model. Consequently, we extended our metabolic model by incorporating diacylglycerol and triacylglycerol biosynthesis pathway (TRIGLSYN-PWY) reactions (EC: 2.3.1.15, 2.3.1.51, 2.3.1.23, 3.1.3.4, 2.3.1.20) for TAG (12:0/14:0/16:0) and TAG (16:1/18:1/18:2) for modeling storage of different fatty acids in BSF larvae biomass.

### Approximation of food intake and transport reaction constraints

*H. illucens* larvae received 8 kg of diet in 9 days composed of a total of 846 g glucose, 1870 g egg white protein with amino acid composition listed in Table 4 and 5284 g of water (Table 3). No glucose was left in the substrate at the end of the 9 days, but there was protein and water residue which was measured during wet lab experiments. We assumed linear amino acid and glucose consumption during the 9 days of larval growth (Table 3).

**Table 3.** Applied diet ingredients, ratios and residue.

| Ingredients | % | g/100g | Total diet, g | Left in residue, g |
| --- | --- | --- | --- | --- |
| Glucose | 10 | 10.58 | 846 | 0 |
| Egg White Protein | 20 | 23.38 | 1870 | 1310 |
| Water | 70 | 66.05 | 5284 | 4890 |

**Table 4.** Dietary amino acid profile conversion to transport reaction rates per 1 larva a day (detailed measurement data in supplementary file 8).

| Amino acid Profile | mg per 100 g protein | Consumed protein weight, g | mg/larva/day | mmol/larva/day |
| --- | --- | --- | --- | --- |
| Lysine | 6000 | 85.272 | 1.769 | 0.012 |
| Methionine | 3500 | 50.116 | 1.039 | 0.007 |
| Cystine | 2500 | 36.465 | 0.756 | 0.003 |
| Aspartate | 4800 | 48.62 | 1.008 | 0.008 |
| Asparagine | 4800 | 48.62 | 1.008 | 0.008 |
| Threonine | 4300 | 62.832 | 1.303 | 0.011 |
| Serine | 6400 | 93.687 | 1.943 | 0.018 |
| Glutamate | 6150 | 62.271 | 1.292 | 0.009 |
| Glutamine | 6150 | 62.271 | 1.292 | 0.009 |
| Proline | 3600 | 52.921 | 1.098 | 0.010 |
| Glycine | 3300 | 47.872 | 0.993 | 0.013 |
| Alanine | 5800 | 84.898 | 1.761 | 0.020 |
| Valine | 6500 | 95.183 | 1.974 | 0.017 |
| Isoleucine | 5100 | 74.239 | 1.540 | 0.012 |
| Leucine | 8000 | 117.062 | 2.428 | 0.019 |
| Tyrosine | 3700 | 54.604 | 1.133 | 0.006 |
| Phenylalanine | 5500 | 78.914 | 1.637 | 0.010 |
| Histidine | 2200 | 32.164 | 0.667 | 0.004 |
| Arginine | 5400 | 79.475 | 1.648 | 0.009 |
| Tryptophan | 1500 | 23.001 | 0.477 | 0.002 |

The dietary amino acid profile was converted to transport reaction rates per 1 larva a day (mmol/1 larva/day) considering the amino acid amount left in the residue as shown in our chemical analysis experiments of BSF larva amino acid composition (Table 4) and having the approximate larva weight measured at the last day of experiment which was 0,028 g.

### Data Analysis

To observe and assess resource allocation changes within our metabolic network based on the applied diet, we performed two sets of *in-silico* experiments (i) simulation of biomass growth rate changes for reference state based on our experimental data and under conditions of increased glucose or essential amino acids (EAA); (ii) simulation of different fatty acid (C12:0, C14:0, C16:0, C16:1, C18:0, C:18:1, C18:2) synthesis rate changes with increased glucose consumption and estimation of dietary carbon conversion efficiency (CCE, Formula 1). During the first set of *in-silico* analysis in the MATLAB environment using the Cobra Toolbox FBA method a steady state for our experimental data constraints was confirmed and biomass-specific growth rate was optimized under conditions of doubled glucose or EAA uptake. Objective function during FBA maximization consequently was set to biomass growth reaction.

The second set of experiments involved the use of FVA analysis preserving 90% of maximal DW biomass-specific growth rate per day as described by (60) for fatty acid (C12:0, C14:0, C16:0, C16:1, C18:0, C18:1, C18:2) synthesis rate modeling under doubled glucose uptake conditions. Dietary carbon conversion efficiency was afterwards estimated using Formula 1 and both states (reference and doubled glucose uptake) (Fig. 5.) were compared.

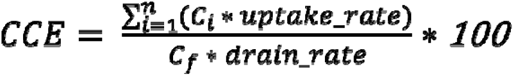

**Formula 1.** Carbon conversion efficiency (CCE) in % estimated using summary diet uptake reaction rates (*uptake_rate*) times carbon atoms per molecule (calculated by FBA in mmol/larva/day) divided by the maximal theoretical fatty acid synthesis rate (*drain_rate*) times carbon atoms per molecule (calculated by FVA in mmol/larva/day) multiplied by 100, where

n - exchange reaction count,

C - carbon atom count in a dietary nutrient (amino acids, glucose, cholesterol),

Cf - carbon atom count in the fatty acid molecule.

## Acknowledgments

FACCE SURPLUS and supported by No. 23-11.17e/20/173 “Sustainable up-cycling of agricultural residues: modular cascading waste conversion system” (UpWaste) and also by the German Federal Ministry of Education and Research (BMBF), in the frame of FACCE-SURPLUS/FACCE-JPI project UpWaste, grant number 031B0934A and 031B0934B. We also thank AGROLAB LUFA for valuable help in the chemical analysis of proteins.

## Data availability

The current manuscript uses *H. illucens* previously published genome annotation data found in (https://www.ncbi.nlm.nih.gov/assembly/GCF_905115235.1).

All experimental measurements, metabolic model and optimization results are included in the article and Supplementary files. All data is available in Git-hub (https://github.com/BigDataInSilicoBiologyGroup/Hermetia_illucens_metabolic_model).. (Supplementary files 2 - 5) All our experimental chemical measurements are available (Supplementary files 8-10). Interactive *H. illucens* larva biochemical network figure manageable in IMFler (Supplementary file 6) and JPEG format (Supplementary file 7).

For additional information and follow-up studies please also visit (https://biomod.lv/).

## References

1. K. Kang, Y. Cai, L. Yue, W. Zhang, Effects of Different Nutritional Conditions on the Growth and Reproduction of Nilaparvata lugens (Stål). Front. Physiol. 12 (2022).

2. Z.-L. Wang, X.-P. Wang, C.-R. Li, Z.-Z. Xia, S.-X. Li, Effect of Dietary Protein and Carbohydrates on Survival and Growth in Larvae of the Henosepilachna vigintioctopunctata (F.) (Coleoptera: Coccinellidae). J. Insect Sci. 18 (2018).

3. K. Berzins, et al., Kinetic and Stoichiometric Modeling-Based Analysis of Docosahexaenoic Acid (DHA) Production Potential by Crypthecodinium cohnii from Glycerol, Glucose and Ethanol. Mar. Drugs 20, 115 (2022).

4. L. Gratani, Plant Phenotypic Plasticity in Response to Environmental Factors. Adv. Bot. 2014, 1–17 (2014).

5. E. M. Sundermann, M. J. Lercher, D. Heckmann, Modeling photosynthetic resource allocation connects physiology with evolutionary environments. Sci. Rep. 11, 15979 (2021).

6. K. B. Barragan-Fonseca, M. Dicke, J. J. A. van Loon, Influence of larval density and dietary nutrient concentration on performance, body protein, and fat contents of black soldier fly larvae (Hermetia illucens). Entomol. Exp. Appl. 166, 761–770 (2018).

7. M. Y. Abduh, M. P. Perdana, M. A. Bara, L. W. Anggraeni, R. E. Putra, Effects of aeration rate and feed on growth, productivity and nutrient composition of black soldier fly (Hermetia illucens L.) larvae. J. Asia. Pac. Entomol. 25, 101902 (2022).

8. T. M. Fowles, C. Nansen, “Insect-Based Bioconversion: Value from Food Waste” in Food Waste Management, (Springer International Publishing, 2020), pp. 321–346.

9. B. Hoc, et al., About lipid metabolism in Hermetia illucens (L. 1758): on the origin of fatty acids in prepupae. Sci. Rep. 10, 11916 (2020).

10. D. Nestel, et al., Resource allocation and compensation during development in holometabolous insects. J. Insect Physiol. 95, 78–88 (2016).

11. M. Filipiak, M. Woyciechowski, M. Czarnoleski, Stoichiometric niche, nutrient partitioning and resource allocation in a solitary bee are sex-specific and phosphorous is allocated mainly to the cocoon. Sci. Rep. 11, 652 (2021).

12. J. M. Monk, et al., iML1515, a knowledgebase that computes Escherichia coli traits. Nat. Biotechnol. 35, 904–908 (2017).

13. H. Lu, et al., A consensus S. cerevisiae metabolic model Yeast8 and its ecosystem for comprehensively probing cellular metabolism. Nat. Commun. 10, 3586 (2019).

14. E. Stalidzans, A. Seiman, K. Peebo, V. Komasilovs, A. Pentjuss, Model-based metabolism design: constraints for kinetic and stoichiometric models. Biochem. Soc. Trans. 46, 261–267 (2018).

15. A. Pentjuss, et al., Biotechnological potential of respiring Zymomonas mobilis: A stoichiometric analysis of its central metabolism. J. Biotechnol. 165, 1–10 (2013).

16. A. Pentjuss, et al., Model-based biotechnological potential analysis of Kluyveromyces marxianus central metabolism. J. Ind. Microbiol. Biotechnol. 44, 1177–1190 (2017).

17. A. Mardinoglu, J. Boren, U. Smith, M. Uhlen, J. Nielsen, Systems biology in hepatology: approaches and applications. Nat. Rev. Gastroenterol. Hepatol. 15, 365–377 (2018).

18. V. Komasilovs, A. Pentjuss, A. Elsts, E. Stalidzans, Total enzyme activity constraint and homeostatic constraint impact on the optimization potential of a kinetic model. Biosystems 162, 128–134 (2017).

19. A. Elsts, A. Pentjuss, E. Stalidzans, SpaceScanner: COPASI wrapper for automated management of global stochastic optimization experiments. Bioinformatics 33, 2966–2967 (2017).

20. S. Choi, C. W. Song, J. H. Shin, S. Y. Lee, Biorefineries for the production of top building block chemicals and their derivatives. Metab. Eng. 28, 223–239 (2015).

21. U. Kalnenieks, A. Pentjuss, R. Rutkis, E. Stalidzans, D. A. Fell, Modeling of Zymomonas mobilis central metabolism for novel metabolic engineering strategies. Front. Microbiol. 5(2014).

22. U. Kalnenieks, et al., Improvement of Acetaldehyde Production in Zymomonas mobilis by Engineering of Its Aerobic Metabolism. Front. Microbiol. 10(2019).

23. C. P. McNally, E. Borenstein, Metabolic model-based analysis of the emergence of bacterial cross-feeding via extensive gene loss. BMC Syst. Biol. 12, 69 (2018).

24. D. Liang, et al., Use of high-resolution metabolomics for the identification of metabolic signals associated with traffic-related air pollution. Environ. Int. 120, 145–154 (2018).

25. C. C. Rittschof, S. Schirmeier, Insect models of central nervous system energy metabolism and its links to behavior. Glia 66, 1160–1175 (2018).

26. N. Y. D. Ankrah, B. Chouaia, A. E. Douglas, The Cost of Metabolic Interactions in Symbioses between Insects and Bacteria with Reduced Genomes. MBio 9(2018).

27. R. J. Hall, S. Thorpe, G. H. Thomas, A. J. Wood, Simulating the evolutionary trajectories of metabolic pathways for insect symbionts in the genus Sodalis. Microb. Genomics 6(2020).

28. C. I. Rumbos, C. G. Athanassiou, ‘Insects as Food and Feed: If You Can’t Beat Them, Eat Them!’—To the Magnificent Seven and Beyond. J. Insect Sci. 21(2021).

29. R. P. Mensink, “Fatty Acids: Health Effects of Saturated Fatty Acids” in Encyclopedia of Human Nutrition, (Elsevier, 2013), pp. 215–219.

30. P. W. Siri-Tarino, Q. Sun, F. B. Hu, R. M. Krauss, Meta-analysis of prospective cohort studies evaluating the association of saturated fat with cardiovascular disease. Am. J. Clin. Nutr. 91, 535–546 (2010).

31. R. Chowdhury, et al., Association of Dietary, Circulating, and Supplement Fatty Acids With Coronary Risk. Ann. Intern. Med. 160, 398 (2014).

32. S. Hamley, The effect of replacing saturated fat with mostly n-6 polyunsaturated fat on coronary heart disease: a meta-analysis of randomised controlled trials. Nutr. J. 16, 30 (2017).

33. B. S. Nurul-Iman, Y. Kamisah, K. Jaarin, H. M. S. Qodriyah, Virgin Coconut Oil Prevents Blood Pressure Elevation and Improves Endothelial Functions in Rats Fed with Repeatedly Heated Palm Oil. Evidence-Based Complement. Altern. Med. 2013, 1–7 (2013).

34. H. Dabadie, E. Peuchant, M. Bernard, P. Leruyet, F. Mendy, Moderate intake of myristic acid in sn-2 position has beneficial lipidic effects and enhances DHA of cholesteryl esters in an interventional study. J. Nutr. Biochem. 16, 375–382 (2005).

35. Y. Kim, J. Lee, S. Park, S. Kim, J. Lee, Inhibition of polymicrobial biofilm formation by saw palmetto oil, lauric acid and myristic acid. Microb. Biotechnol. 15, 590–602 (2022).

36. J. A. Menendez, L. Vellon, R. Colomer, R. Lupu, Oleic acid, the main monounsaturated fatty acid of olive oil, suppresses Her-2/neu (erbB-2) expression and synergistically enhances the growth inhibitory effects of trastuzumab (Herceptin^TM^) in breast cancer cells with Her-2/neu oncogene amplification. Ann. Oncol. 16, 359–371 (2005).

37. J. Menendez, R. Lupu, Mediterranean Dietary Traditions for the Molecular Treatment of Human Cancer: Anti-Oncogenic Actions of the Main Olive Oils Monounsaturated Fatty Acid Oleic Acid (18:1n-9). Curr. Pharm. Biotechnol. 7, 495–502 (2006).

38. L. Schwingshackl, G. Hoffmann, Monounsaturated fatty acids, olive oil and health status: a systematic review and meta-analysis of cohort studies. Lipids Health Dis. 13, 154 (2014).

39. J. M. Bourre, “Brain lipids and ageing” in Food for the Ageing Population, (Elsevier, 2009), pp. 219–251.

40. N. H. Choulis, “Miscellaneous drugs, materials, medical devices, and techniques” in (2011), pp. 1009–1029.

41. S. Mukerjee, A. S. Saeedan, M. N. Ansari, M. Singh, Polyunsaturated Fatty Acids Mediated Regulation of Membrane Biochemistry and Tumor Cell Membrane Integrity. Membranes (Basel). 11, 479 (2021).

42. J. W. Pelley, “Fatty Acid and Triglyceride Metabolism” in Elsevier’s Integrated Review Biochemistry, (Elsevier, 2012), pp. 81–88.

43. T. Brody, “VITAMINS” in Nutritional Biochemistry, (Elsevier, 1999), pp. 491–692.

44. T. A. B. Sanders, “Introduction” in Functional Dietary Lipids, (Elsevier, 2016), pp. 1–20.

45. T. Spranghers, et al., Nutritional composition of black soldier fly (Hermetia illucens) prepupae reared on different organic waste substrates. J. Sci. Food Agric. 97, 2594–2600 (2017).

46. B. A. Rumpold, O. K. Schlüter, Potential and challenges of insects as an innovative source for food and feed production. Innov. Food Sci. Emerg. Technol. 17, 1–11 (2013).

47. N. Ewald, et al., Fatty acid composition of black soldier fly larvae (Hermetia illucens) – Possibilities and limitations for modification through diet. Waste Manag. 102, 40–47 (2020).

48. S. Kahwaji, M. B. Johnson, A. C. Kheirabadi, D. Groulx, M. A. White, Fatty acids and related phase change materials for reliable thermal energy storage at moderate temperatures. Sol. Energy Mater. Sol. Cells 167, 109–120 (2017).

49. X. Li, et al., Growth and Fatty Acid Composition of Black Soldier Fly Hermetia illucens (Diptera: Stratiomyidae) Larvae Are Influenced by Dietary Fat Sources and Levels. Animals 12, 486 (2022).

50. A. M. Feist, B. O. Palsson, The biomass objective function. Curr. Opin. Microbiol. 13, 344–349 (2010).

51. C. Thiele, J. Spandl, Cell biology of lipid droplets. Curr. Opin. Cell Biol. 20, 378–385 (2008).

52. D. Murphy, The biogenesis and functions of lipid bodies in animals, plants and microorganisms. Prog. Lipid Res. 40, 325–438 (2001).

53. M. A. Welte, A. P. Gould, Lipid droplet functions beyond energy storage. Biochim. Biophys. Acta - Mol. Cell Biol. Lipids 1862, 1260–1272 (2017).

54. A.-M. Reimers, A. C. Reimers, The steady-state assumption in oscillating and growing systems. J. Theor. Biol. 406, 176–186 (2016).

55. J. D. Orth, I. Thiele, B. Ø. Palsson, What is flux balance analysis? Nat. Biotechnol. 28, 245–248 (2010).

56. J. D. Orth, B. Ø. Palsson, R. M. T. Fleming, Reconstruction and Use of Microbial Metabolic Networks: the Core Escherichia coli Metabolic Model as an Educational Guide. EcoSal Plus 4(2010).

57. J. M. Monk, et al., Genome-scale metabolic reconstructions of multiple Escherichia coli strains highlight strain-specific adaptations to nutritional environments. Proc. Natl. Acad. Sci. 110, 20338–20343 (2013).

58. E. Brunk, et al., Recon3D enables a three-dimensional view of gene variation in human metabolism. Nat. Biotechnol. 36, 272–281 (2018).

59. S. Gudmundsson, I. Thiele, Computationally efficient flux variability analysis. BMC Bioinformatics 11, 489 (2010).

60. M. M. Seyedalmoosavi, M. Mielenz, T. Veldkamp, G. Daş, C. C. Metges, Growth efficiency, intestinal biology, and nutrient utilization and requirements of black soldier fly (Hermetia illucens) larvae compared to monogastric livestock species: a review. J. Anim. Sci. Biotechnol. 13, 31 (2022).

61. J. W. Schönborn, L. Jehrke, T. Mettler-Altmann, M. Beller, FlySilico: Flux balance modeling of Drosophila larval growth and resource allocation. Sci. Rep. 9, 17156 (2019).

62. L. Broeckx, et al., Growth of Black Soldier Fly Larvae Reared on Organic Side-Streams. Sustainability 13, 12953 (2021).

63. A. van Huis, D. G. A. B. Oonincx, The environmental sustainability of insects as food and feed. A review. Agron. Sustain. Dev. 37, 43 (2017).

64. D. Zhou, J. Yuan, Y. Zhou, Y. Liu, Preparation and characterization of myristic acid/expanded graphite composite phase change materials for thermal energy storage. Sci. Rep. 10, 10889 (2020).

65. M. Ataman, D. F. Hernandez Gardiol, G. Fengos, V. Hatzimanikatis, redGEM: Systematic reduction and analysis of genome-scale metabolic reconstructions for development of consistent core metabolic models. PLOS Comput. Biol. 13, e1005444 (2017).

66. R. Petrovs, E. Stalidzans, A. Pentjuss, IMFLer: A Web Application for Interactive Metabolic Flux Analysis and Visualization. J. Comput. Biol. 28, 1021–1032 (2021).

67. A. Giannetto, et al., Hermetia illucens (Diptera: Stratiomydae) larvae and prepupae: Biomass production, fatty acid profile and expression of key genes involved in lipid metabolism. J. Biotechnol. 307, 44–54 (2020).

68. I. Opatovsky, T. Vitenberg, A. Jonas-Levi, R. Gutman, Does Consumption of Baker’s Yeast (Saccharomyces cerevisiae) by Black Soldier Fly (Diptera: Stratiomyidae) Larvae Affect Their Fatty Acid Composition? J. Insect Sci. 21(2021).

69. S. Smetana, M. Palanisamy, A. Mathys, V. Heinz, Sustainability of insect use for feed and food: Life Cycle Assessment perspective. J. Clean. Prod. 137, 741–751 (2016).

70. S. Smetana, E. Schmitt, A. Mathys, Sustainable use of Hermetia illucens insect biomass for feed and food: Attributional and consequential life cycle assessment. Resour. Conserv. Recycl. 144, 285–296 (2019).

71. J. Cammack, J. Tomberlin, The Impact of Diet Protein and Carbohydrate on Select Life-History Traits of The Black Soldier Fly Hermetia illucens (L.) (Diptera: Stratiomyidae). Insects 8, 56 (2017).

72. A. Gligorescu, S. Toft, H. Hauggaard-Nielsen, J. A. Axelsen, S. A. Nielsen, Development, metabolism and nutrient composition of black soldier fly larvae (Hermetia illucens_; Diptera: Stratiomyidae) in relation to temperature and diet. J. Insects as Food Feed 4, 123–133 (2018).

73. C. D. Miranda, J. A. Cammack, J. K. Tomberlin, Mass Production of the Black Soldier Fly, Hermetia illucens (L.), (Diptera: Stratiomyidae) Reared on Three Manure Types. Animals 10, 1243 (2020).

74. S. Bellezza Oddon, I. Biasato, A. Resconi, L. Gasco, Determination of lipid requirements in black soldier fly through semi-purified diets. Sci. Rep. 12, 10922 (2022).

75. H. Wang, et al., RAVEN 2.0: A versatile toolbox for metabolic network reconstruction and a case study on Streptomyces coelicolor. PLOS Comput. Biol. 14, e1006541 (2018).

76. P. D. Karp, et al., The BioCyc collection of microbial genomes and metabolic pathways. Brief. Bioinform. 20, 1085–1093 (2019).

77. M. Kanehisa, M. Furumichi, M. Tanabe, Y. Sato, K. Morishima, KEGG: new perspectives on genomes, pathways, diseases and drugs. Nucleic Acids Res. 45, D353–D361 (2017).

78. K. Degtyarenko, et al., ChEBI: a database and ontology for chemical entities of biological interest. Nucleic Acids Res. 36, D344–D350 (2007).

79. J. D. Orth, R. M. T. Fleming, B. Ø. Palsson, Reconstruction and Use of Microbial Metabolic Networks: the Core Escherichia coli Metabolic Model as an Educational Guide. EcoSal Plus 4(2010).

80. L. Heirendt, et al., Creation and analysis of biochemical constraint-based models using the COBRA Toolbox v.3.0. Nat. Protoc. 14, 639–702 (2019).

81. C. Ceccotti, et al., New value from food and industrial wastes – Bioaccumulation of omega-3 fatty acids from an oleaginous microbial biomass paired with a brewery by-product using black soldier fly (Hermetia illucens) larvae. Waste Manag. 143, 95–104 (2022).

