## Supplementary material for "Metabolic modeling of *Hermetia illucens* larvae resource allocation for high-value fatty acid production": Supplemetary materials: 1_Supplemental_information..docx

1. File of Supplemental table of content.
2. *H. illucens* medium scale metabolic model in JSON file format (Compatible with CobraPy)
3. *H. illucens* medium scale metabolic model in MATLAB Mat file format (Compatible with Cobra Toolbox 3.0)
4. *H. illucens* medium scale metabolic model in XLSX file format (Compatible with Cobra Toolbox 3.0 with xlsx import support)
5. *H. illucens* medium scale metabolic model in SBML (XML) file format compatible with other metabolic modelling tools.
6. *H. illucens* medium scale metabolic model interactive biochemical pathways interactive map, which is compatible with IMFler interactive online modelling tool.
7. *H. illucens* medium scale metabolic model interactive biochemical pathways map in JPG file format
8. *H. illucens* Larva and diet composition experimental measurements
9. *H. illucens* Larva calculated biomass function measurements
10. *H. illucens* Chemcial analysis experimentals measurements for larva and diet composition (preliminary data)
