## Supplementary figures and images for "Metabolic modeling of *Hermetia illucens* larvae resource allocation for high-value fatty acid production"

### 7_Hermetia_network_Fig_5.jpg

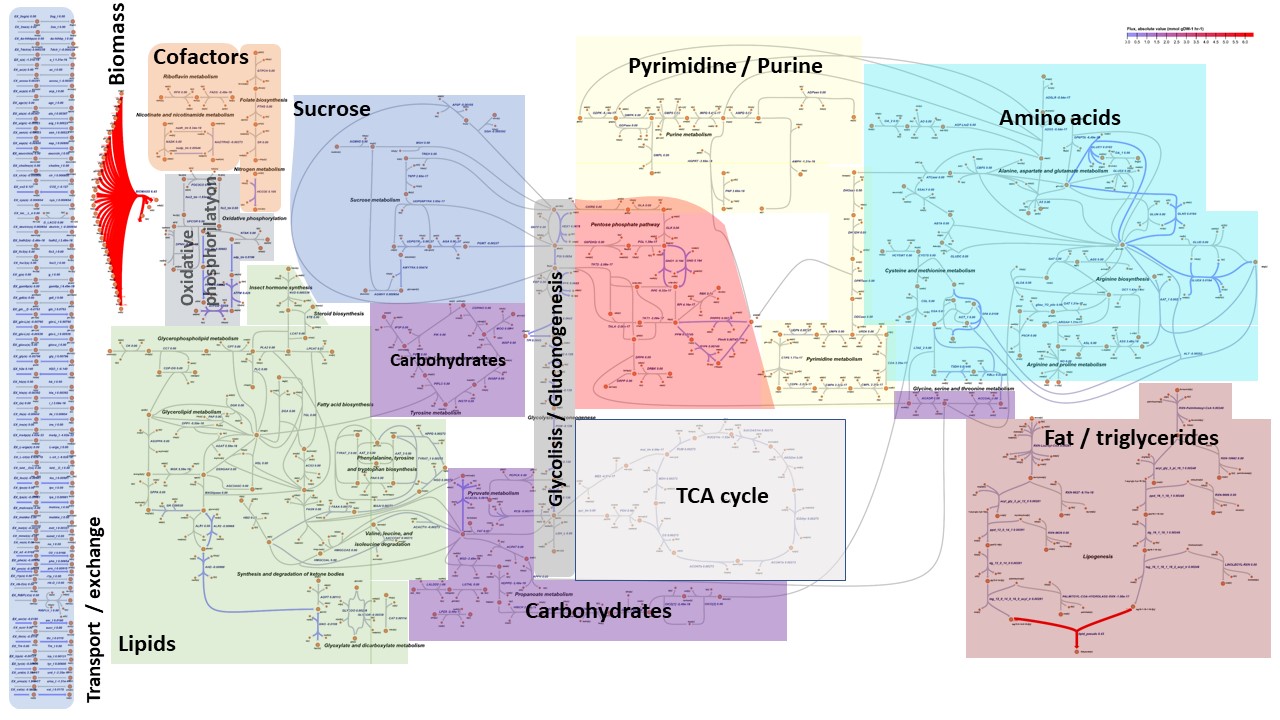
